# Guanine crystals discovered in bacteria

**DOI:** 10.1101/2022.12.02.518751

**Authors:** María Elisa Pavan, Federico Movilla, Esteban E. Pavan, Florencia Di Salvo, Nancy I. López, M. Julia Pettinari

**Affiliations:** Departamento de Química Biológica, Facultad de Ciencias Exactas y Naturales, Universidad de Buenos Aires, Buenos Aires, Argentina; Departamento de Química Inorgánica, Analítica y Química Física, CONICET-Instituto de Química Física de los Materiales, Medio Ambiente y Energía (INQUIMAE), Facultad de Ciencias Exactas y Naturales, Universidad de Buenos Aires, Buenos Aires, Argentina; Biomedical Technologies Laboratory, Department of Electronics, Information and Bioengineering, Politecnico di Milano, Milan, Italy; IQUIBICEN-CONICET, Facultad de Ciencias Exactas y Naturales, Universidad de Buenos Aires, Buenos Aires, Argentina

**Keywords:** biogenic guanine crystals, guanine monohydrate, melanin, *Aeromonas*, biomaterial, bacterial guanine crystals

## Abstract

Guanine crystals are organic biogenic crystals found in many organisms. Due to their exceptionally high refractive index, they contribute to structural color and are responsible for the reflective effect in the skin and visual organs in animals such as fish, reptiles and spiders. Occurrence of these crystals in animals has been known for many years, and they have also been observed in eukaryotic microorganisms, but not in prokaryotes. In this work we report the discovery of extracellular crystals in bacteria, and reveal that they are composed of guanine, and particularly the unusual monohydrate form. We demonstrate the occurrence of these crystals in *Aeromonas* and other bacteria, and investigate the metabolic traits related to their synthesis. In all cases studied the presence of the guanine crystals in bacteria correlate with the absence of guanine deaminase, which could lead to guanine accumulation providing the substrate for crystal formation. Our finding of the hitherto unknown guanine crystal occurrence in prokaryotes extends the range of guanine crystal producing organisms to a new domain of life. Bacteria constitute a new and more accessible model to study the process of guanine crystal formation and assembly. This discovery opens countless chemical and biological questions, including those about the functional and adaptive significance of their production in these microorganisms. It also paves the road for the development of simple and convenient processes to obtain biogenic guanine crystals for diverse applications.

**Significance:** Guanine crystal formation is well known in animals such as fish, reptiles and arthropods (among other eukaryotic organisms), but its occurrence has never been reported in prokaryotes. This manuscript describes the discovery of extracellular guanine crystals in bacteria, and reveals that they are composed of the unusual monohydrate form of guanine. Knowledge of guanine crystal biosynthesis in bacteria could lead to a better understanding of their synthesis in other organisms. It also paves the road for the development of simple and convenient processes to obtain biogenic guanine crystals for diverse applications. Our finding extends the range of guanine crystal producing organisms to a new domain of life.

## Main

Guanine is a purine, one of the four bases of the nucleotides that constitute the backbone of nucleic acids. Guanine crystals have been observed in diverse organisms^1,2,3^. The most widely studied are related to the production of structural color or are part of the reflective tissue in visual organs in many animals, including arthropods, mollusks, amphibians, reptiles and fish^2^. The extensive occurrence of guanine crystals in optical systems is probably due to its exceptionally high refractive index, and to the fact that guanine is a widespread and abundant metabolite^4^. Guanine can also be excreted as an end product of nitrogen metabolism. Guanine crystals have long been known to be among the main excretion products in arachnids^5^, and more recently found in land crustaceans^6^. In eukaryotic microorganisms, guanine crystal-like particles were observed in the cytoplasm of paramecia and other protozoa^7^, and in several microalgae^8^ such as, dinoflagellates^9^. Guanine crystals in these organisms have been proposed to act as purine storage reservoirs, formed through the excretion of purine excess, and used as a source of purines and organic nitrogen during starvation^7,8^.

There are three crystal forms for guanine, two polymorphs of the anhydrous phases, the α and β forms^10,11^, and the monohydrate^12^. The three forms have been obtained *in vitro*, displaying different morphologies: the α and/or β polymorphs showed a prismatic bulky morphology while guanine monohydrate formed elongated needle-like crystals^13^. Biogenic guanine crystals are typically composed of anhydrous guanine, and studies that have analyzed its crystalline form (obtained from spiders, fish and copepods) confirmed the presence of the β polymorph in all cases^13^.

While guanine crystals have been observed in diverse groups of animals and in eukaryotic microorganisms, these biogenic crystals have not been reported in prokaryotes. Analysis of bright crystals observed in colonies of melanogenic *Aeromonas salmonicida* subsp. *pectinolytica* 34mel^T^^14^ revealed that the crystals are composed of guanine, and particularly the unusual monohydrate form. Careful examination allowed the discovery of guanine crystals in other bacteria as well. This work describes the characteristics of bacterial guanine crystals and investigates the metabolic traits that could lead to the synthesis of these crystals in bacteria.

## Results and discussion

### Crystals found in bacterial colonies

Serendipitous observation of month-old colonies of the melanogenic bacterium *A. salmonicida* subsp. *pectinolytica* strain 34mel^T^ (from now on, 34mel) revealed the presence of glimmering crystals in contrast with the dark background. These particles were associated to the colonies and not the surrounding medium and appeared as birefringent crystalline material under polarized light (*SI Appendix*, Fig. S1). Scanning Electron Microscopy (SEM) showed that the crystalline material consisted of mesoscopically structured 50 to 100 μm sphere-like aggregates of elongated nanocrystals (Fig. 1a-c and *SI Appendix*, Fig. S2c). The crystals were also observed when 34mel was cultured in liquid medium (Fig. 1d-f). Organization of biogenic crystals in complex mesoscopic structures such as the skeletal structures composed of calcium carbonate found in sea urchin spines and mollusc nacre have been extensively studied^15^. Organic biogenic crystals such as those composed of guanine often have special arrangements that can potentiate their properties^16^. Guanine crystals found in animals are normally described as platelets or prisms that can be arranged in blocks and are often found in specific layered tissues^1, 2,17^. In eukaryotic microorganisms packed prismatic particles of guanine crystals have been observed inside intracellular vesicles^3,18^. The rounded aggregates of nanocrystals produced by the bacteria are very different from the structures observed in other biogenic crystals.

**Fig. 1.**
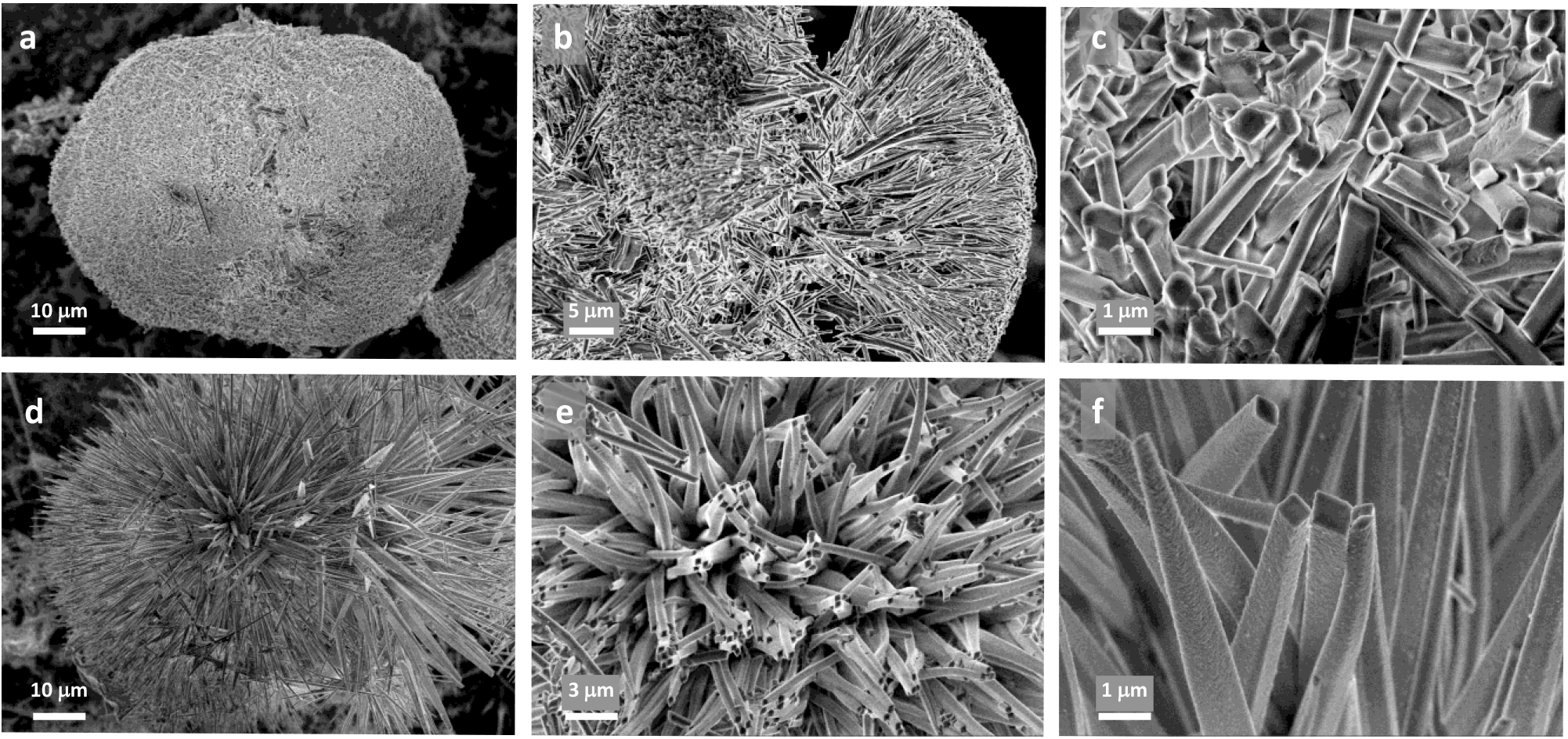
SEM micrographs of the crystalline material produced by 34mel. **a-f,** Detail of crystalline aggregates and individual nanocrystals when grown in solid (**a-c**) and liquid (**d-f**) LB medium.

The morphology of the nanocrystals found in 34mel can be described predominantly as rhomboidal or hexagonal elongated prisms, with an average size for the base of around 500 nm x 350 nm, and a maximum length of about 10 μm (Fig. 1 and *SI Appendix*, Fig. S2). The size of the base of the nanocrystals is comparable to prismatic biogenic guanine crystals observed in some spiders^17^ but the bacterial crystals are considerably more elongated (Fig. 1).

### Characterization of the crystalline material

The structural analysis of the crystals produced by 34mel was performed on bulky samples obtained after collection, washing and drying under vacuum, using different spectroscopy and X-ray diffraction studies (XRD) and CHN elemental analysis. High-resolution electrospray ionization mass spectroscopy (HR ESI-MS) of the crystals showed a signal at m/z 152.0574 corresponding to the ion [M+H]^+^ (Fig. 2a and *SI Appendix*, Fig. S3). MS/MS experiments for the target ion gave place to the expected fragments for guanine^19^ (Fig. 2b). The assignment was confirmed by comparison with commercial guanine run under the same experimental conditions (*SI Appendix*, Fig. S4). The aggregation of guanine in solution is clearly demonstrated by the presence of ions with m/z > 152 associated to [*n*M + H]^+^ and [*n*M + Na]^+^ mainly (*SI Appendix*, Fig. S3). ^1^H NMR (*SI Appendix*, Fig. S5), FT-IR (Fig. 2d), UV-visible (*SI Appendix*, Fig. S6) spectroscopy and XRD (Fig. 2c and *SI Appendix*, Fig. S7) results confirmed that the crystalline material corresponded to guanine crystals.

**Fig. 2.**
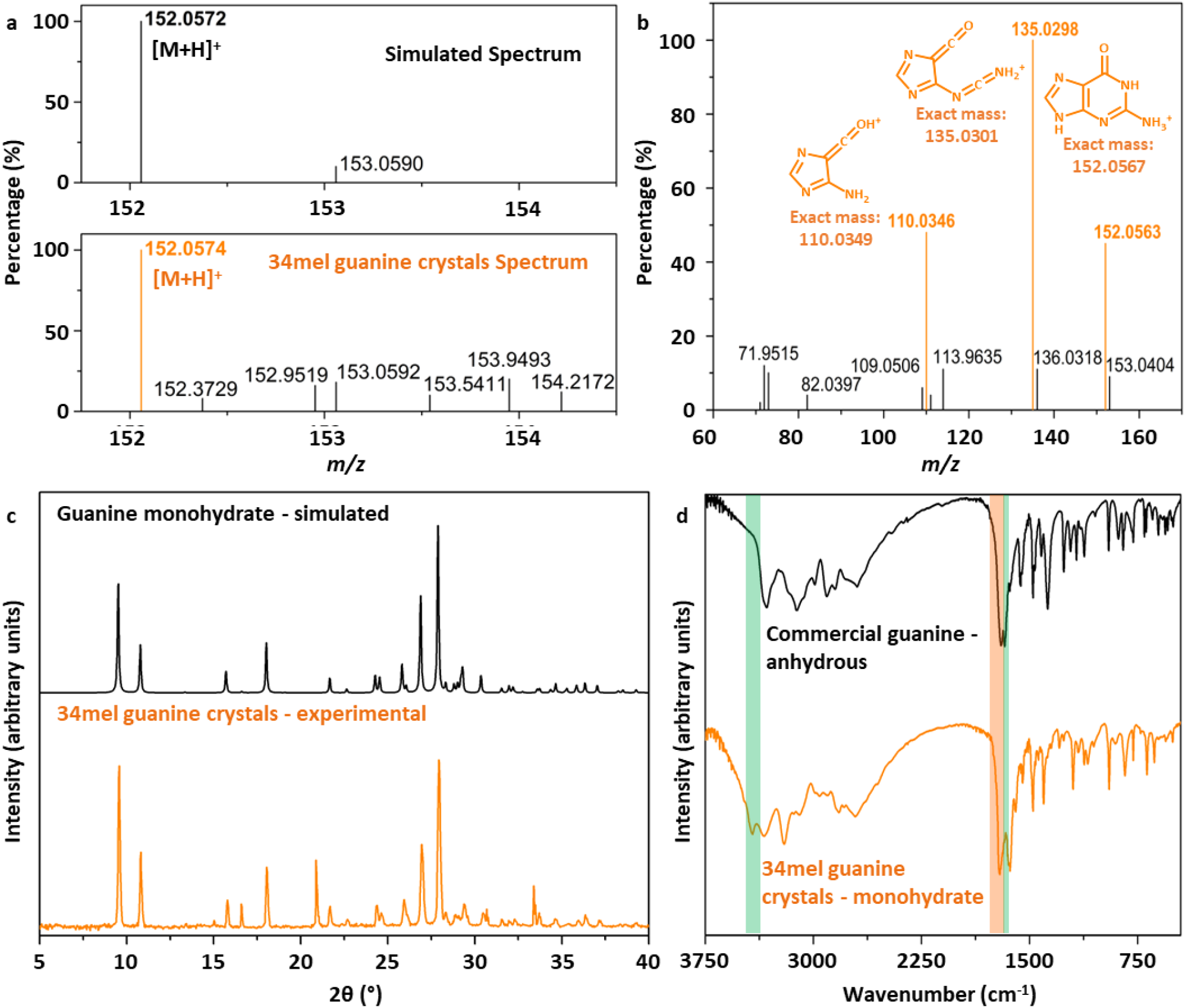
Characterization of guanine crystals produced by 34mel. **a,** ESI-MS spectrum of biogenic guanine compared to the simulated data for guanine. **b,** ESI MS/MS spectrum of the [M + H]^+^ ion m/z 152.0574. Solvent: methanol: H_2_O. In color, proposed structures for MS/MS obtained ions are shown. **c,** Powder X-ray diffraction pattern of the biogenic guanine crystals and the simulated from single-crystal X-ray diffraction data for the guanine monohydrate phase^12^. **d,** FT-IR spectra of biogenic guanine crystals and commercial guanine, signals associated with water molecules are highlighted in green and those of carbonyl and amine groups in orange.

Although these characterization techniques allowed us to determine that guanine was the major component in all the crystalline samples, only solid state characterization studies gave information regarding the crystalline form of the guanine produced by 34mel. Powder X-ray diffraction patterns of biogenic guanine samples showed a very good fit with the data reported for the monohydrate phase, one of the three crystal forms of guanine known to date^10–12^ (Fig. 2c and *SI Appendix*, Fig. S7). In the FT-IR spectrum, the signals at 3425 and 3200 cm^-1^, and the one at 1590 cm^-1^, associated to the stretching modes ν_1_ and ν_3_, and the bending mode ν_2_ of water molecules, respectively, and the carbonyl and primary amine stretching modes that generate two resolved signals at 1681 cm^-1^ and 1632 cm^-1^, are also in agreement with previously reported data for the guanine monohydrate crystalline phase (Fig. 2d)^13^. Elemental analysis revealed a high N content compound (C 35.2%, H 4.2%, N 37.1%), similar to the calculated composition for a sample of guanine monohydrate considering traces of melanin and water (see *SI Appendix* for details).

Guanine monohydrate crystals are very hard to produce in the laboratory. Their crystal structure was determined in 1971^12^ and only a few years ago a detailed study of guanine crystallization in solution provided experimental data of this phase^13^. As described, guanine crystals found in 34mel are elongated prisms (Fig. 1b,c,e,f), a crystalline habit resembling the one of the guanine monohydrate crystals obtained *in vitro*^13^ and different from the crystals formed by anhydrous guanine^2^.

Guanine crystals produced by 34mel are brown even after several steps of purification with different solvents. This coloring could be associated with the homogentisate melanin synthesized by the microorganism if traces of the pigment were included in the crystalline structure of the guanine. Crystallization experiments of commercial guanine were performed *in vitro*, including melanin (obtained from the bacteria) in the solution, and the crystalline material obtained was analyzed through powder XRD (see *SI Appendix*, for details). For every condition tested, the same crystalline phase was observed for the samples with or without biogenic melanin (*SI Appendix*, Figs. S8 and S9). Furthermore, guaninium chloride dihydrate crystalline material obtained from solutions containing biogenic melanin was suitable for single crystal XRD structural determination. Crystallographic data did not show any dye molecules in the structure (*SI Appendix*, Fig. S10 and Table S1). These results confirmed that the color providing substance (melanin), did not alter the crystal packing and structural parameters (*SI Appendix*, Table S2). A recent study showed that intracrystalline dopants do not alter the morphology of biogenic guanine crystals^17^.

### Biological and genetic aspects of guanine crystal synthesis

Guanine crystals are found in many animals in specialized cells such as guanocytes in spiders^20^ or iridophores in fish^21^. In eukaryotic microorganisms including many microalgae, intracellular guanine crystals are found in vacuoles^3,18^. The formation of intracellular crystals in bacteria has been reported in very few cases, such as the membrane surrounded magnetite crystals in magnetotactic bacteria^22^, or the parasporal crystals formed by the Cry protein in *Bacillus thuringiensis*^23^. The bacterial guanine crystals observed in this study are extracellular, with a size that is several times larger than the cells, and structured in large crystalline aggregates, differing both in location and morphology when compared to biogenic crystals formed by other organisms.

In nature, purines are synthesized as nucleotides in the so-called *de novo* pathway. Enzymatic removal of the phosphate and sugar yields the corresponding bases. Nucleosides and free bases released from nucleic acid breakdown are recycled through the salvage pathway, or degraded to uric acid. Guanine accumulation leading to crystal formation in animals has been related to upregulation of the guanine portion of the *de novo* purine synthesis^24^ or attributed to deficiencies in enzymes involved in guanine degradation, such as xanthine dehydrogenase^5^ or guanine deaminase^25,26^.

Purine metabolic pathways were analyzed in 34mel, all sequenced *Aeromonas* and some related bacteria. Special emphasis was placed on guanine degradation and the purine nucleotide salvage pathway (Fig. 3a).

**Fig. 3.**
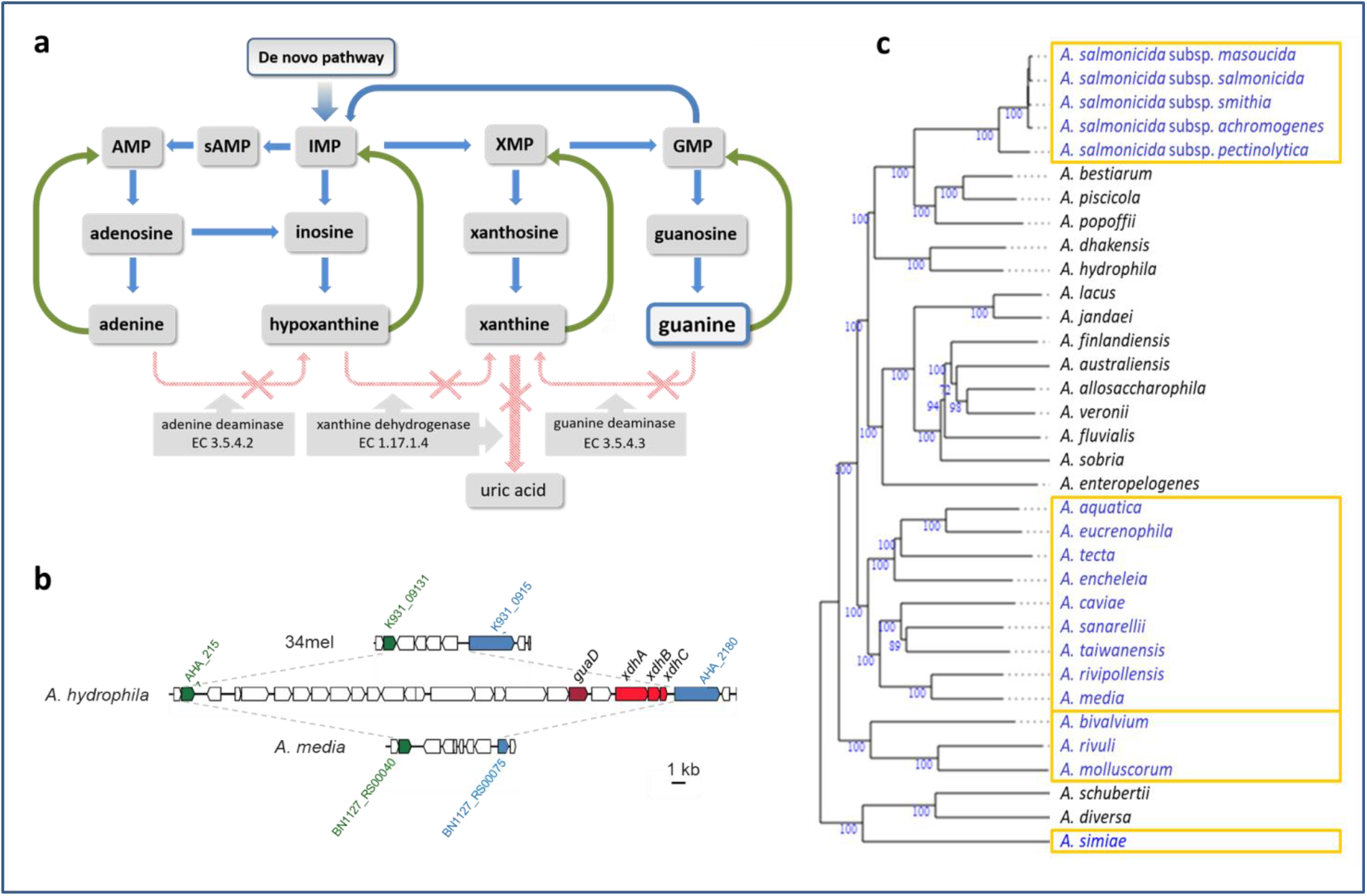
Purine metabolism in *Aeromonas*. **a,** Purine metabolic pathway in 34mel. *De novo* formation of purines (blue arrows) and salvage reactions (green arrows). Enzymes absent in 34mel with the corresponding E.C. numbers are shown with crossed out arrows. **b,** Comparison of the genomic region containing *guaD* and *xdhABC* in *A. hydrophila* showing their absence in representative *Aeromonas*. Homologous flanking genes are shown: Auxin efflux carrier family transporter (green) and ExeM/NucH family extracellular endonuclease (blue) along with the locus tags in each genome. **c,** Phylogenomic tree of *Aeromonas* species showing the presence (black) or the absence (blue) of the gene encoding the guanine deaminase. The numbers below branches are GBDP pseudo-bootstrap support values > 60 % from 100 replications, with an average branch support of 94.4%. The tree was rooted at the midpoint^27^. The strains used and the accession numbers of their genomes are indicated in *SI Appendix*, Table S3.

Interestingly, the gene coding for the guanine deaminase, commonly present in prokaryotes^28^, is absent in 34mel. The lack of this enzyme would prevent the degradation of guanine that could only be recycled back to the nucleoside monophosphate through the salvage pathway (Fig. 3a), potentially leading to guanine excess that would be available for crystal formation. A comparative genomic search in all *Aeromonas* revealed that the genes coding for the guanine deaminase and the three xanthine dehydrogenase subunits are clustered in some species such as *A. hydrophila*. However, this gene cluster is absent in 34mel (Fig 3b) and in about half of the genomes, and a deeper analysis suggested the occurrence of deletions that affect this region, involving different phylogenetic clades (Fig. 3c). Alterations in the salvage pathway can also lead to guanine accumulation. For example, an *E. coli* strain accumulates guanine due to a mutation in *gpt*, the gene encoding guanine phosphoribosyltransferase, the enzyme that catalyzes the conversion of guanine to GMP in the guanine salvage^29^. This *E. coli* strain was grown in LB to analyze if guanine accumulation led to crystal formation, but no crystals were observed after more than 30 days.

Since 34mel lacks the guanine deaminase and produces melanin, the possible relationship of melanin synthesis with guanine crystal formation was investigated by analyzing these traits and the occurrence of crystals in *Aeromonas* with different combinations (Table 1). Guanine crystals were observed in melanogenic *A. media* (Fig. 4d and *SI Appendix*, Fig. S2) and *A. salmonicida* subsp. *salmonicida*, but also in the non melanogenic *A. salmonicida* subsp. *masoucida* (Fig. 4b) and *A. caviae*. All these guanine crystal forming bacteria are devoid of guanine deaminase (Table 1). In contrast, no crystals were produced by *A. hydrophila* that carries the guanine deaminase. Careful observation of cultures of field strains of *A. allosaccharophila, A. bestiarum*, or *A. veronii*^30^ showed that they were also devoid of guanine crystals. Although there is no available sequence information for the strains used, analysis of the genomes of sequenced type strains of these species has revealed the presence of the genes coding for the guanine deaminase (Table 1).

**Fig. 4.**
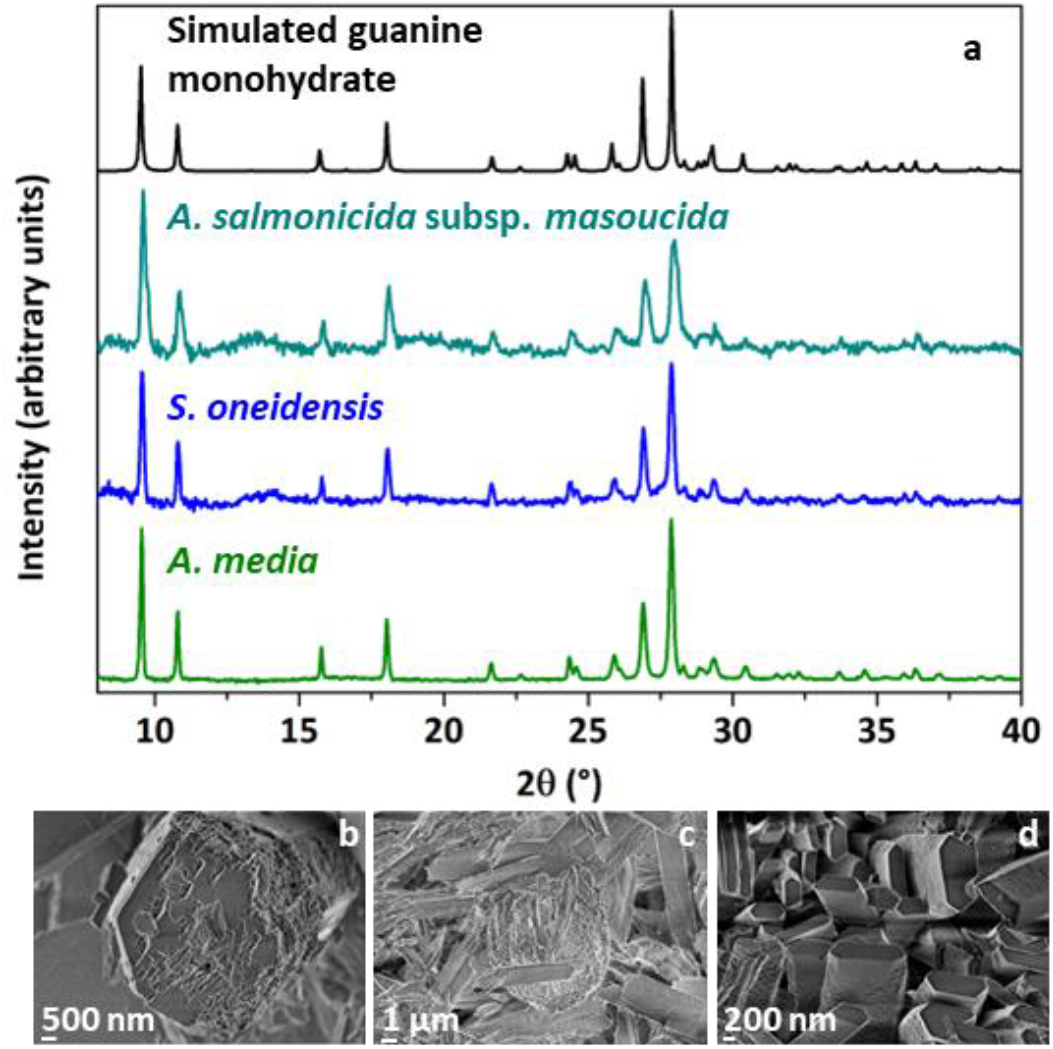
Characterization of guanine crystals produced by several bacteria. **a,** Comparison of powder X-ray diffraction profiles for guanine crystals produced by the different bacteria and the simulated pattern from single crystal X-ray diffraction data for guanine monohydrate phase^12^. **b-d**, SEM micrographs of the guanine crystals produced by the bacteria *A. salmonicida* subsp. *masoucida* (**b**), *S. oneidensis* **(c**) and *A. media* (**d**).

**Table 1.** Experimental analysis of guanine crystal formation and melanin production in selected bacterial species, along with presence or absence of genes encoding guanine deaminase (*guaD*) and xanthine dehydrogenase (*xdh*) in the genomes.

|  | Genes |  | Phenotype |  |
| --- | --- | --- | --- | --- |
|  | *guaD* | *xdh* | Melanin | Guanine crystals |
| *A. salmonicida* subsp. *pectinolytica* 34mel<sup>T</sup> | no | no | ● | ◇ |
| *A. salmonicida* subsp. *salmonicida* ATCC 33658<sup>T</sup> | no | no | ● | ◇ |
| *A. salmonicida* subsp. *masoucida* NBRC 13784<sup>T</sup> | no | no | - | ◇ |
| *A. media* CECT 4232<sup>T</sup> | no | no | ● | ◇ |
| *A. hydrophila* ATCC 7966<sup>T</sup> | yes | yes | - | - |
| *A. caviae* C | no* | no* | - | ◇ |
| *A. allosaccharophila* A1 | yes* | yes* | - | - |
| *A. bestiarum* B1 | yes* | yes* | - | - |
| *A. veronii* V1 | yes* | yes* | - | - |
| *S. oneidensis* MR-1<sup>T</sup> | no | no | ● | ◇ |
| *E. coli* K-12 substr. BW25113 | yes | yes | - | - |
| *P. aeruginosa* PAO1 | yes | yes | - | - |
| *P. extremaustralis* DSM 25547 | yes | yes | - | - |
| *P. protegens* Pf-5 | yes | yes | - | - |
| *P. putida* KT2440 | yes | yes | - | - |
| *P. syringae* B728a | yes | yes | - | - |

\*Genome for this strain is not available, but the gene(s) is (are) present or absent in the type strain of the species (*S*) Appendix, Table S3). Presence of melanin and/or guanine crystals is indicated by colored circles or diamonds.

When cultures of other bacteria were searched for guanine crystals it was observed that *E. coli* and several species of *Pseudomonas* did not produce them, and analysis of their genomes revealed that they carry the guanine deaminase gene (Table 1). Guanine crystals were found in *Shewanella oneidensis*, a melanin producing bacterium that lacks the guanine deaminase gene (as observed in crystal producing *Aeromonas*) (Table 1). Crystals formed in melanogenic *A. media* and *S. oneidensis* and in non-melanogenic *A. salmonicida* subsp. *masoucida* observed through SEM (Fig. 4b-d) have different sizes but share a prismatic crystal morphology. Powder XRD studies revealed that the diffraction patterns of the crystalline material found in these bacteria have a very good agreement with the calculated data for the guanine monohydrate crystal form (Fig. 4a). These results suggest that this composition could be characteristic of bacterial guanine crystals, differing from the composition commonly found in eukaryotes. All reports of guanine crystals in animals have described them as composed of anhydrous guanine^11^. In the case of eukaryotic microorganisms, although purine crystals have been known for many years, their composition has been investigated in detail in the last years^3^. In a very recent study that investigated crystalline inclusions in diverse unicellular eukaryotes, almost all guanine crystals contained anhydrous guanine with just a few examples containing the monohydrate form^31^.

Guanine crystals are associated with melanin in many organisms^2,32,33^. In bacteria that produce homogentisate melanin such as 34mel, melanin synthesis can be inhibited by the herbicide bicyclopyrone^34^. When 34mel was grown in the presence of this inhibitor crystals with similar morphology were observed, although delayed by several days (*SI Appendix*, Fig. S2). The occurrence of guanine crystals in non melanogenic bacteria, together with the observation of crystals in 34mel in the presence of the inhibitor, indicate that melanin synthesis is not essential for guanine crystal formation.

The results presented in this work show that the presence of the crystals in bacteria correlated with the absence of guanine deaminase, which could lead to guanine accumulation providing the substrate for crystal formation. Furthermore, a phylogenetic analysis of the occurrence of deletions involving the gene coding for this enzyme within the genus *Aeromonas* revealed that its loss seems to be the result of several independent events (Fig. 3c). When guanine crystal production was studied in animals, a patchy phylogenetic distribution was observed^2^, suggesting that both in bacteria and in animals, guanine crystal formation arose several times independently.

The existence of guanine crystals in several groups of animals has been known for many years, and their contribution to structural color and as part of reflective tissues has been extensively studied^2,35^. Their occurrence in other organisms, such as unicellular eukaryotes, was studied many years ago^9,36^, and thought to be limited to a few cases. Recent renewed interest in guanine crystals and the application of new technologies has expanded this knowledge, and a very recent study showed them to be widespread among eukaryotic microorganisms^31^. Our work has demonstrated their presence in prokaryotes, extending the range of guanine crystal producing organisms to a new Domain of life. Future studies will indicate if this capability is restricted to a few bacteria or is more extended among the prokaryotes. The finding of the hitherto unknown guanine crystal formation in prokaryotes has opened countless chemical and biological questions, including those about the functional and adaptive significance of their production in these microorganisms.

## Materials and methods

### Bacterial strains and culture conditions

*A. salmonicida* subsp. *pectinolytica* 34mel^T^ (DSM 12609^T^), *A. salmonicida* subsp. *masoucida* NBRC 13784^T^, *A. media* CECT 4232^T^, *A. hydrophila* ATCC 7966^T^ and field strains *A. caviae* C, *A. allosaccharophila* A1, *A. bestiarum* B1 and *A. veronii* V1^30^ were grown at 28°C. *A. salmonicida* subsp. *salmonicida* ATCC 33658^T^ was grown at 24°C. *Escherichia coli* K-12 substr. BW25113, *Shewanella oneidensis* MR-1^T^, *Pseudomonas aeruginosa* PAO1, *Pseudomonas extremaustralis* DSM 25547, *Pseudomonas protegens* Pf-5, *Pseudomonas putida* KT2440 and *Pseudomonas syringae* pv. *syringae* B728a were grown at 28°C. All strains were grown in lysogeny broth (LB) medium except for *S. oneidensis*, that was grown in tryptic soy agar (TSA). After 5 days incubation cultures were kept at room temperature or 4°C and crystals formation was followed using an Olympus Tokyo CK inverted microscope or a stereoscopic microscope Nikon SMZ-745T. For melanin synthesis inhibition 1 mM bicyclopyrone was added to the growth medium^34^.

### Characterization of biogenic guanine crystals

Crystals were collected from solid or liquid cultures washing out bacteria and culture residues with water (for SEM) or with solvents with decreasing polarity (water, ethanol and acetone, for XRD, NMR, and ESI-MS experiments), with gentle agitation. Solvent residue was removed by vacuum drying. The crystalline material was then characterized using different techniques (polarized light microscopy, SEM, UV-vis, FT-IR, ESI-MS & MS/MS and ^1^H NMR).

Light micrographs using polarized Light Microscopy (PLM) were taken with a stereoscopic trinocular microscope Nikon SMZ-745T that includes a lighting system Nikon Ni-150. Images were processed using the programs Micrometrics^™^ SE Premium and ImageJ^37^. SEM images were produced using a Carl Zeiss NTS – SUPRA 40. UV–visible spectra of guanine crystals in acid solution (HCl pH 2) were recorded using a Hewlett-Packard 8453 diode array spectrometer. Elemental analysis was carried out in a Carlo Erba CHNS EA-1108 microanalyzer using atropine as standard. FT-IR spectra were recorded using a Nicolet Avatar 320 FTIR spectrometer with a Spectra Tech cell for KBr pellets. High-resolution electrospray ionization mass spectroscopy (HR ESI-MS) was performed using crystals dissolved in a mixture of methanol: DMSO or methanol: H_2_O. Mass spectra were recorded on a Xevo G2S Q-TOF (Waters Corp.) instrument, using an electrospray ionization source and quadrupole-flight time analyzer in methanol: water 80:20 or DMSO as solvent. ^1^H-NMR spectra were recorded using a Bruker AM500 equipped with a broadband probe. ^1^H shifts are reported relative to DMSO-*d*6 (*δ*) 2.50 ppm.

### Powder X-ray diffraction (powder XRD)

Data were recorded on a PANalytical Empyrean diffractometer equipped with a 4-kW sealed tube Cu Kα X-ray radiation (generator power settings: 60 kV and 100 mA) and a PIXcel^3D^ area detector using parallel beam geometry (1/2-1-8mm slits, 15mm incident mask). Samples were packed on a silicon monocrystal sample holder that was then placed on the sample holder attachment. For all pXRD experiments the data were collected over an angle range 5° to 50° with a scanning speed of 23 s per step with 0.026° step.

### Comparative genome analysis

The genomes of *Aeromonas* strains and other *Gammaproteobacteria* used for comparative analysis are shown in *SI Appendix*, Table S3. Global alignments of the genome of 34mel with those of bacterial strains belonging to the genus *Aeromonas* or *Shewanella* were performed using the Needleman-Wunsch algorithm (using the BLOSUM50 scoring matrix and a maximum gap open penalty of 10), which is included in the Bioinformatics Toolbox of Matlab^38^. Phylogenomic tree of *Aeromonas* species was constructed using the tools included in Type Strain Genome server (TYGS), with FastME 2.1.6.1^39^ from GBDP distances calculated from genome sequences. The branch lengths are scaled in terms of GBDP distance formula d5.

## Data availability

All data generated or analyzed during this study are included in this published article (and its supplementary information files). Crystallographic data for guaninium chloride dihydrate - melanin have been deposited at the Cambridge Crystallographic Data Centre (CCDC) under the deposition number 2156488.

## ACKNOWLEDGMENTS

Roberto Servant and Dr Laura Levin for encouraging M.E.P. during the first steps of this research. Alejandro Perretta, Cynthia Sequeiros and Jorge Trelles, who kindly provided us with some *Aeromonas* strains. We gratefully acknowledge UBA (329 20020170100310BA, 20020170100433BA) and ANPCYT (PICT 2016-621) for funding resources. FDS, NIL and MJP are staff members of CONICET. FM acknowledges the Universidad de Buenos Aires (UBA) for his scholarships.

## Author contributions

M.E.P. made the original discovery. M.E.P., N.I.L. and M.J.P. conceived the project and performed microbiological experiments and genomic analysis. M.E.P., N.I.L., M.J.P., F.D.S. and F.M. carried out crystal collection and SEM analysis. F.D.S. and F.M. performed chemical characterization experiments, spectroscopic and XRD data analyses. E.E.P. conducted the bioinformatics analyses and genomic comparisons. All authors discussed the results and commented on the manuscript. All authors have given approval to the final version of the manuscript.

