## Supporting information for "Guanine crystals discovered in bacteria"

Author affiliations

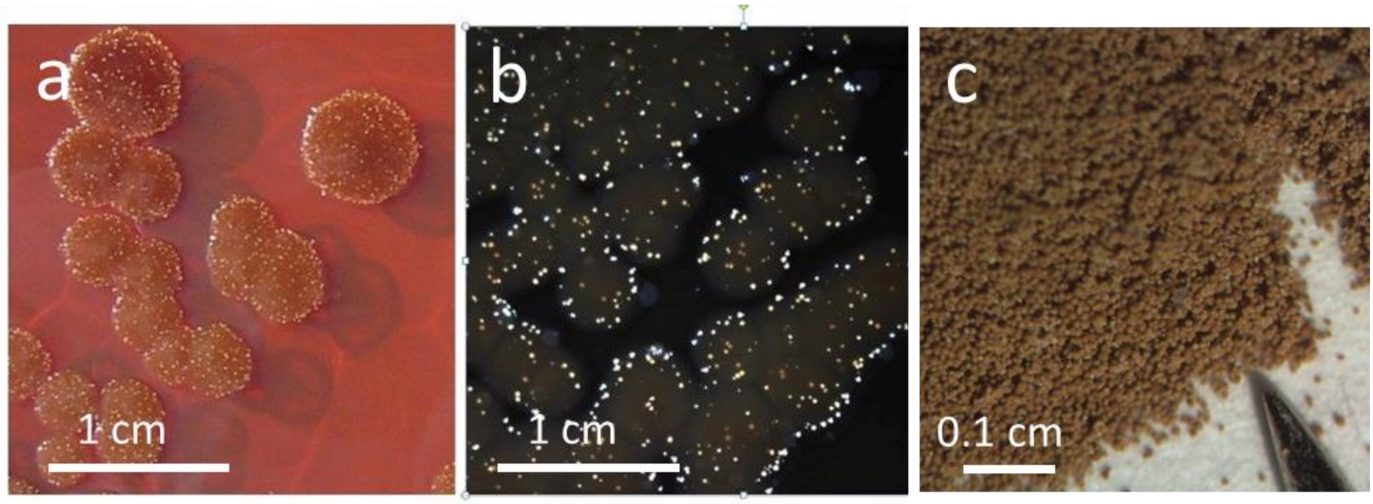


**Fig. S1. Crystalline aggregates in colonies from a month-old plate of *A. salmonicida* subsp. *pectinolytica* 34mel. a,** Bacterial colonies displaying the brown color due to melanin production. The crystal aggregates are seen as bright particles. **b,** Bacterial colonies seen under polarized light microscopy revealing the birefringence of the crystalline aggregates. **c,** Guanine crystals collected from several plates. The tip of a common pin is shown for size comparison.

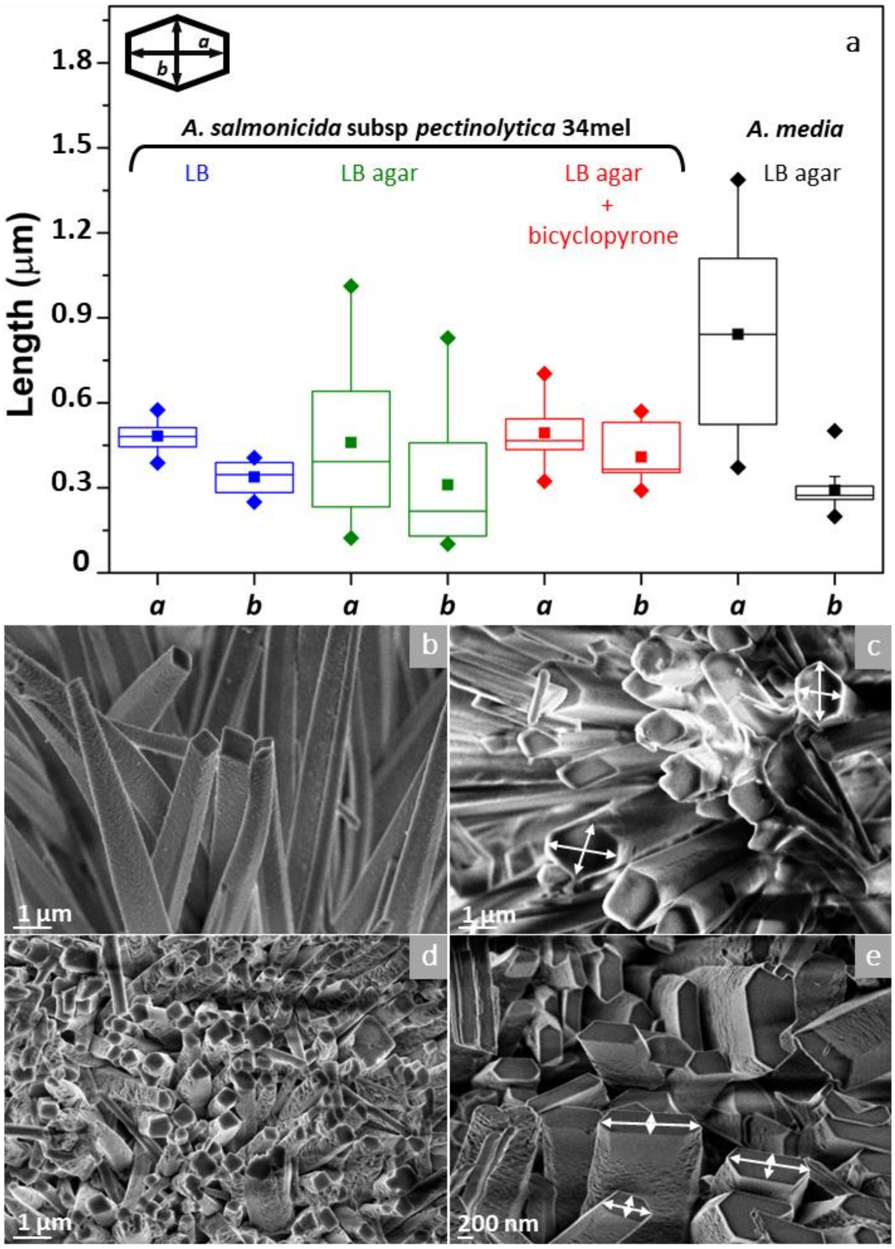


**Fig. S2. Size distribution of the base of the prismatic crystals formed by 34mel grown in different conditions and by *A. media*. a,** The base of individual crystals was measured for each experimental condition (n=12) and data analyzed using ImageJ[37].The box limits represent the range between the first and third quartiles for each condition, the center lines show the median, the square symbols represent the mean, and the ends of the whiskers extend to 1.5× the interquartile range. The diagram in the top left corner shows the long (a) and short (b) axis used for size analysis. **b-e**, SEM micrographs of the biogenic crystals of 34mel grown on liquid culture (**b**), agar plates (**c**), agar plates with the inhibitor bicyclopyrone (**d**) and *A. media* in agar plates (**e**).

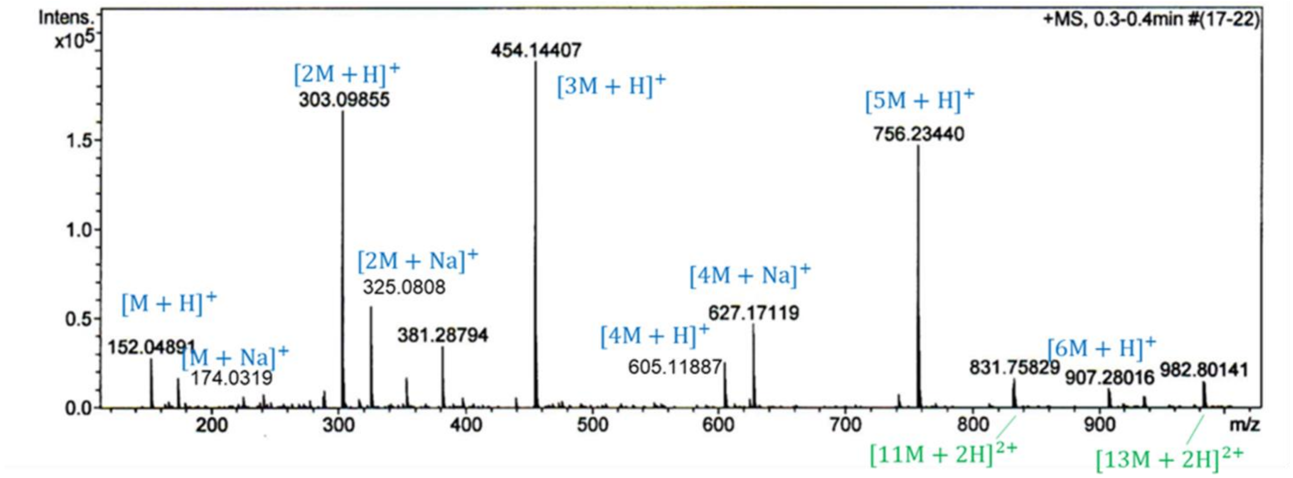


**Fig. S3. ESI-MS experiment for guanine produced by 34mel.** The spectrum shows guanine association in solution. Solvent: methanol: DMSO 80:20.

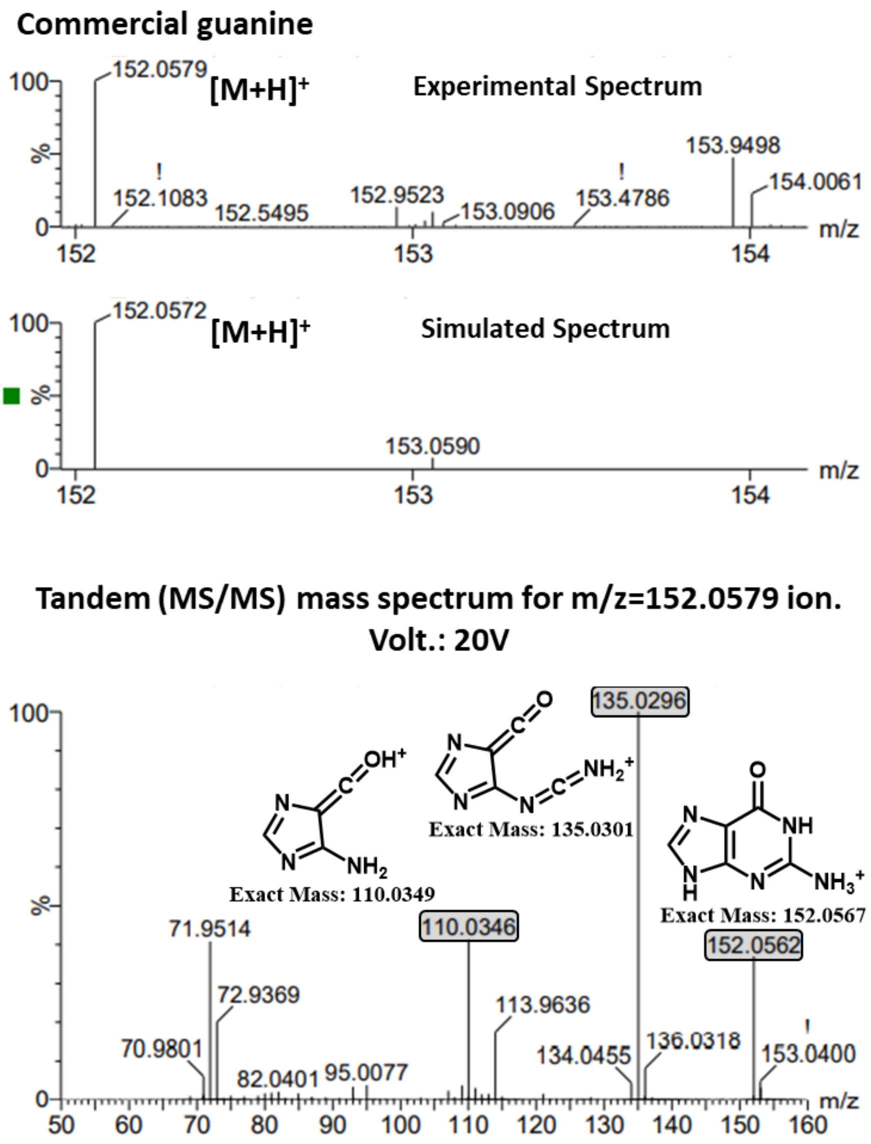


**Fig. S4. ESI-MS and MS/MS experiments for commercial guanine. a**, ESI-MS experiment; $[M+H]^+$ ion (m/z 152.0579) is indicated. **b**, MS/MS experiment using $[M+H]^+$ ion (m/z 152.0579) as the parent ion. Solvent: methanol: $H_2O$.

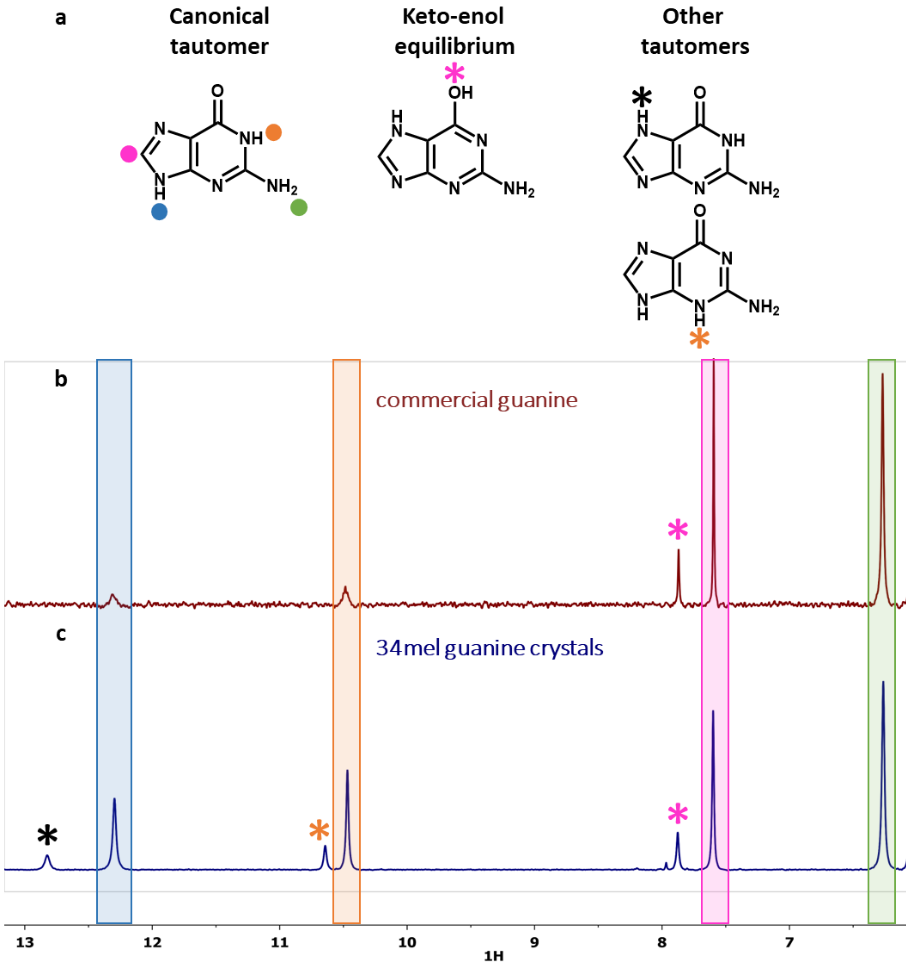


**Fig. S5. [1]H-NMR experiments. a**, Scheme of some of the expected tautomeric forms of guanine. **b** and **c**, [1]H-NMR spectra for commercial guanine (**b**) and guanine produced by 34mel (**c**). In color, signal assignment for guanine hydrogens and proposed signal assignment that may be produced from tautomers in solution.

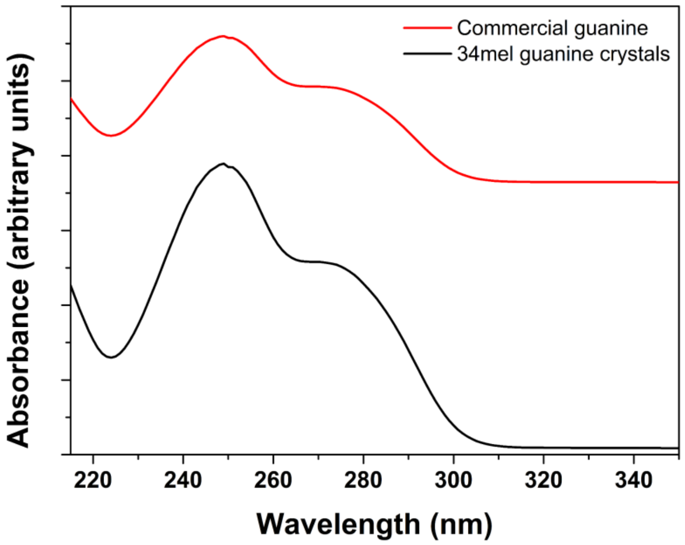


**Fig. S6. UV-vis spectra for commercial and biogenic guanine in acid solution**.

### Elemental analysis

The presence of biogenic melanin and water molecules were taken into account in the calculated values of C% H% and N% for the guanine samples. Biogenic melanin was represented using a pyomelanin monomer which corresponds to a homogentisic acid unit with two hydrogen atoms removed [$(C_8H_4O_4)_n$]. A good agreement was obtained when considering 6% of the monomer and 14% of water [$(C_5H_5N_5O)_{15}(C_8H_4O_4).22H_2O$].

### Crystallization Experiments and Additional X-ray Diffraction Studies

#### Synthesis of the guanine crystalline phases

**Anhydrous β-phase.** Crystals were produced by adding 20 mg of guanine powder (Sigma−Aldrich) in 600 μL of pure water and then, the solid was completely dissolved after dropwise addition of a solution of NaOH 1M. The crystallization of the desired phase was induced by adjusting the pH of the solution to a value of 10. The resulting clear solution was left open to the air at room temperature until white crystalline solids were observed. Crystalline material was then separated by centrifugation, purified by several additions of distilled water and subsequent centrifugation and finally, dried under vacuum.

**Anhydrous β-phase with melanin.** Crystals were produced following the same procedure as described previously but, instead of pure water, filtered melanized liquid culture was used to prepare the starting solution. As a result, a brown crystalline solid was obtained.

**Anhydrous α-phase.** Crystals were produced by adding 20 mg of guanine powder (Sigma−Aldrich) in 600 μL of pure water and then, the solid was partially dissolved after dropwise addition of a solution of HCl 1M. The crystallization of the desired phase was induced by adjusting the pH of the solution to a value of 2. The resulting suspension was sonicated for 5 minutes and then heated at 90°C for an hour, as a result a clear solution was obtained. Crystalline material was then separated by centrifugation, purified by several additions of distilled water and subsequent centrifugation and finally, dried under vacuum.

**Anhydrous α-phase with melanin.** Crystals were produced following the same procedure as described previously but, instead of pure water, filtered melanized liquid culture was used to prepare the starting solution. Due to the low pH, biogenic melanin remained insoluble and guanine was not fully dissolved after sonication and heating thus, the resultant turbid solution was filtered before crystallization. Crystalline material was then separated by centrifugation, purified by several additions of distilled water and subsequent centrifugation and finally, dried under vacuum. As a result, a pale-brown crystalline solid was obtained.

**Guaninium chloride dihydrate.** Crystals were produced by adding 20 mg of guanine powder (Sigma−Aldrich) in 500 µL of pure water and then 200 µL of HCl (c) was added (pH ~ 0). The resultant suspension was warmed until complete dissolution of the guanine. The resulting clear solution was left open to the air at room temperature until white elongated needle-like crystals were observed. Crystalline material was then separated by centrifugation, purified by several additions of distilled water and subsequent centrifugation and finally dried under vacuum.

**Guaninium chloride dihydrate with melanin**. Crystals were produced following the same procedure as described previously but instead of pure water, filtered melanized liquid culture was used to prepare the starting solution. Due to the low pH, biogenic melanin (BM) remained precipitated and guanine was not fully dissolved after sonication and heating thus, the resultant turbid solution was filtered before crystallization. Crystalline material was then separated by centrifugation, purified by several additions of distilled water and subsequent centrifugation and finally dried under vacuum. As a result, pale-brown elongated needle-like crystals were obtained; this material resulted to be suitable for single crystal X-ray diffraction studies.

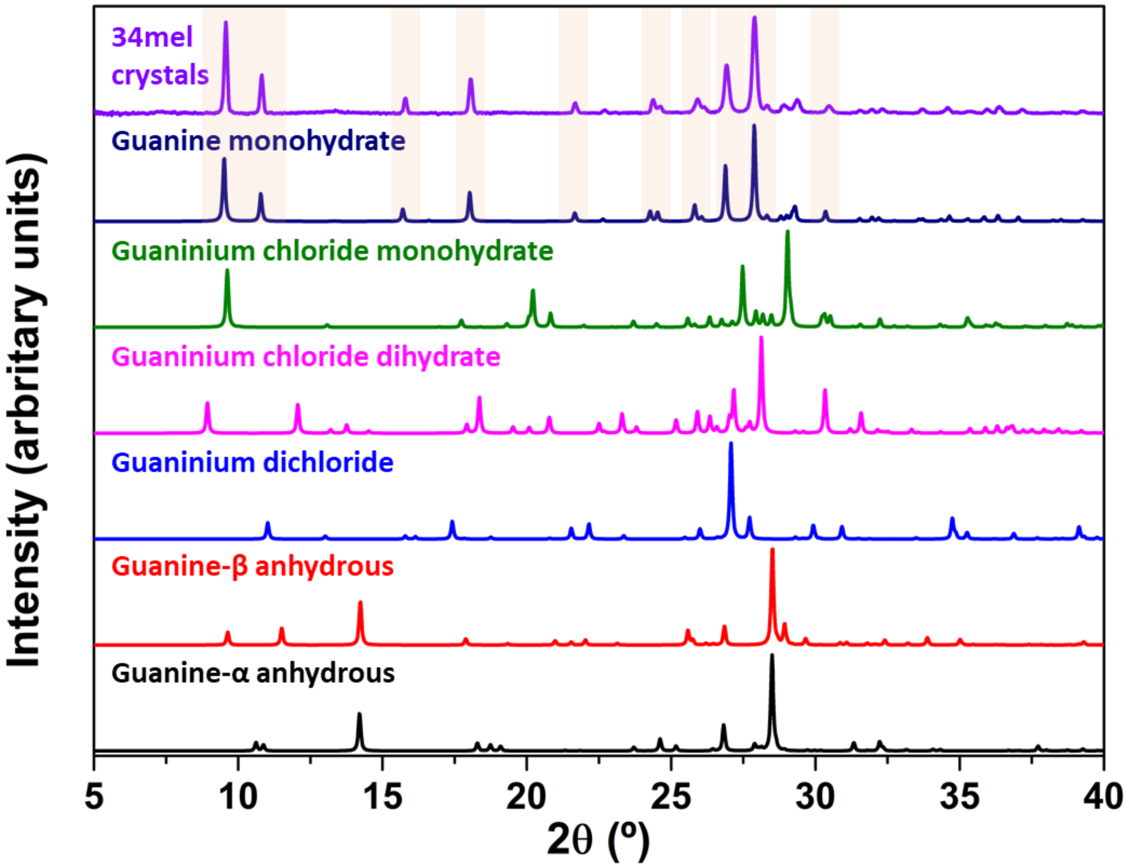


**Fig. S7. X-ray diffraction studies of different crystal forms of guanine**. Experimental powder X-ray diffraction data obtained for guanine crystals produced by 34mel (violet) and simulated patterns from single crystal X-ray diffraction data for guanine monohydrate[12] (dark blue), guaninium chloride monohydrate (green), guaninium chloride dihydrate (pink), guaninium dichloride (blue), β[11] (red) and α[10] (black) anhydrous forms.

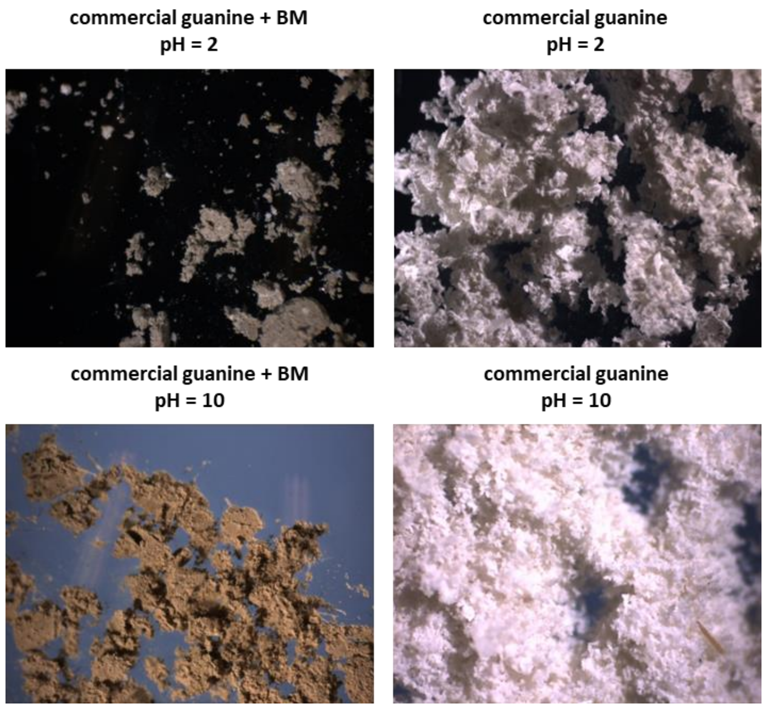


**Fig. S8. Crystalline material obtained as result of crystallization experiments of commercial guanine adding homogentisate melanin synthesized by 34mel (BM) to the solution**. X-ray diffraction patterns of these samples are shown in Fig. S9.

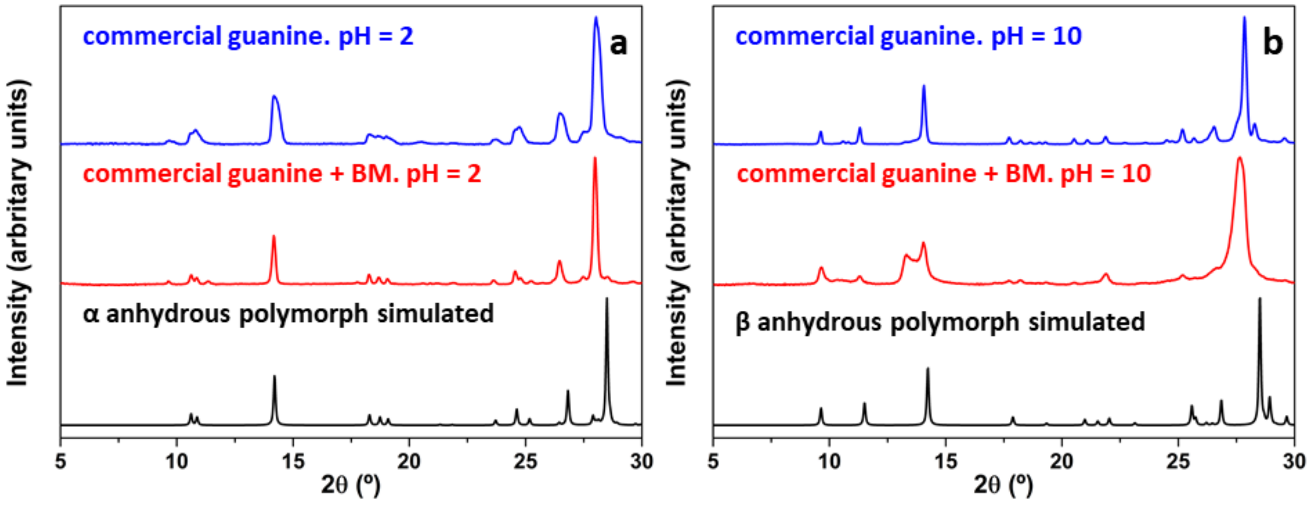


**Fig. S9. Powder X-ray diffraction experiments of the crystalline material obtained under different crystallization conditions**. **a** and **b**, Diffraction patterns at pH 2 (**a**) and pH 10 (**b**) for solutions containing commercial guanine with or without melanin produced by 34mel (BM).

### Single Crystal X-ray diffraction (XRD)

Crystals of guaninium chloride dihydrate crystallized from a solution containing biogenic melanin (BM) were studied by single crystal XRD using an Oxford Diffraction Gemini E lab diffractometer with Mo Kα (λ = 0.71 Å) radiation (Fig. S10). The full data collection was planned using the

CrysAlis Pro strategy tool[40] and data was reduced using CrysAlis Pro software. A Gaussian method implemented in WinGX[41], or a numerical model[42] was used for the absorption correction. Using Olex2[43] the structures were solved by Intrinsic phasing employing ShelXL[44] and refined with the ShelXL[45] package using Least Squares minimization. Non-hydrogen atoms were anisotropically refined. Hydrogen atoms were mostly included at geometrically calculated positions with thermal parameters derived from the parent atoms. Hydrogen atoms attached to the water molecules were located on Fourier maps, fixed, and given isotropic displacement parameters depending on the parent atoms. Chloride anions disorder was modeled over two sets of sites with occupancies of 0.26 and 0.74 (Table S1). Disorder was not modeled for solvent molecules and is represented through large ellipsoids. Crystallographic additional information and tables are included in the ESI. CCDC Deposition Number is 2156488.

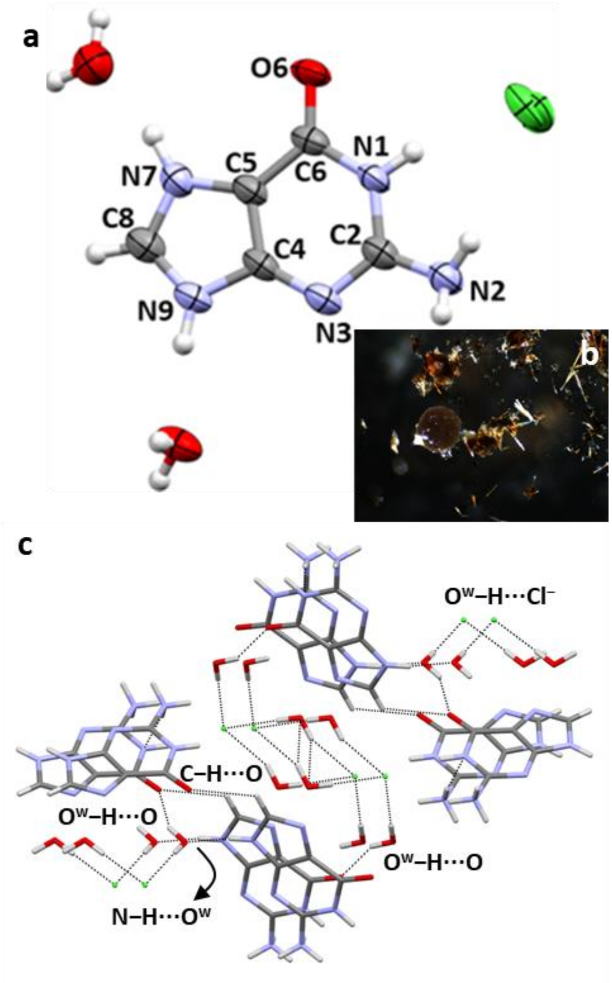


**Fig. S10. Structure of guaninium chloride dihydrate crystallized in presence of melanin produced by the 34mel (BM) determined by single crystal X-ray diffraction**. **a**, ORTEP plot of the asymmetric unit with displacement ellipsoids drawn at 50% probability level, with numbering. **b**, Image of the crystals. **c**, H-bonded supramolecular network.

**Table S1. Crystal data and structure refinement for guaninium chloride dihydrate crystallized in presence of melanin produced by the bacteria (BM).**

| Crystal data and structure refinement for Guaninium chloride dihydrate - BM | |
|---|---|
| Empirical formula | C5H10ClN5O3 |
| Formula weight | 223.63 |
| Temperature/K | 298 |
| Crystal system | monoclinic |
| Space group | P21/c |
| a/Å | 4.8276(10) |
| b/Å | 13.417(2) |
| c/Å | 14.6730(14) |
| α/° | 90 |
| β/° | 94.127(14) |
| γ/° | 90 |
| Volume/Å3 | 947.9(3) |
| Z | 4 |
| ρcalcg/cm3 | 1.567 |
| μ/mm-1 | 0.395 |
| F(000) | 464 |
| Crystal size/mm3 | 0.1 × 0.05 × 0.05 |
| Radiation | MoKα (λ = 0.71073) |
| 2Θ range for data collection/° | 8.242 to 57.544 |
| Index ranges | -5 ≤ h ≤ 6, -14 ≤ k ≤ 18, -19 ≤ l ≤ 18 |
| Reflections collected | 6247 |
| Independent reflections | 2201 [Rint = 0.1620, Rsigma = 0.1534] |
| Data/restraints/parameters | 2201/0/148 |
| Goodness-of-fit on F2 | 0.992 |
| Final R indexes [I>=2σ (I)] | R1 = 0.0907, wR2 = 0.2024 |
| Final R indexes [all data] | R1 = 0.1606, wR2 = 0.2638 |
| Largest diff. peak/hole / e Å-3 | 0.43/-0.49 |

| Bond lengths for Guaninium chloride dihydrate - BM | | |
|---|---|---|
| **Atom** | **Atom** | **Length(Å)** |
| O1 | C2 | 1.231(5) |
| N1 | C1 | 1.370(5) |
| N1 | C2 | 1.389(6) |
| N2 | C1 | 1.336(5) |
| N2 | C3 | 1.334(6) |
| N3 | C4 | 1.373(6) |
| N3 | C5 | 1.309(5) |
| N4 | C3 | 1.382(5) |
| N4 | C5 | 1.328(6) |
| N5 | C1 | 1.321(6) |
| C2 | C4 | 1.420(5) |
| C3 | C4 | 1.381(6) |

| Atomic occupancy for Guaninium chloride dihydrate - BM | |
|---|---|
| **Atom** | ***Occupancy*** |
| Cl1 | 0.78(5) |
| Cl2 | 0.22(5) |

| Hydrogen bonds for Guaninium chloride dihydrate-BM | | | | |
|---|---|---|---|---|
| **D** | **H** | **A** | **d(D-A) (Å)** | **D-H-A (°)** |
| O2 | H2A | O11 | 2.786(5) | 151.9 |
| O3 | H3B | Cl22 | 2.95(4) | 153.3 |
| N4 | H4 | O2 | 2.687(5) | 170.8 |
| N3 | H3 | O3 | 2.645(6) | 169(4) |
| 11+X,3/2-Y,-1/2+Z; 2-1+X,3/2-Y,-1/2+Z | | | | |

**Table S2.** Comparative analysis of different guaninium chloride dihydrate single crystal X-ray diffraction data

| | **a** | **b** | **c** | **α** | **β** | **γ** | **Space Group** | **Ref.** |
|---|---|---|---|---|---|---|---|---|
| **Guaninium chloride dihydrate** (GUANCD01) | 4.8708(7) | 13.237(3) | 14.638(2) | 90 | 93.906 | 90 | P 21/c | [46] |
| **Guaninium chloride dihydrate** (GUANCD02) | 4.8587(11) | 13.228(3) | 14.612(3) | 90 | 93.862(4) | 90 | P 21/c | [47] |
| **Guaninium chloride dihydrate- BM** | 4.8276(10) | 13.417(2) | 14.6730(14) | 90 | 94.127(14) | 90 | P 21/c | *this work* |

**Genome analysis**

**Table S3.** Bacteria used for genome analysis

| **Species** | **GenBank Accession** |
|---|---|
| *Aeromonas allosaccharophila* CECT 4199$^{T}$ | NZ_CDBR00000000 |
| *Aeromonas aquatica* AE235$^{T}$ | NZ_JRGL01000000 |
| *Aeromonas australiensis* CECT 8023$^{T}$ | NZ_CDDH00000000 |
| *Aeromonas bestiarum* CECT 4227$^{T}$ | NZ_CDDA00000000 |
| *Aeromonas bivalvium* CECT 7113$^{T}$ | NZ_CDBT00000000 |
| *Aeromonas caviae* CECT 838$^{T}$ | NZ_JAGDEN000000000.1 |
| *Aeromonas dhakensis* AAK1 | NZ_BAFL00000000 |
| *Aeromonas diversa* CECT 4254$^{T}$ | NZ_CDCE00000000.1 |
| *Aeromonas encheleia* CECT 4342$^{T}$ | NZ_CDDI00000000 |
| *Aeromonas enteropelogenes* CECT 4255$^{T}$ | NZ_CDDE00000000 |
| *Aeromonas eucrenophila* CECT 4224$^{T}$ | NZ_CDDF00000000 |
| *Aeromonas finlandiensis* 4287D$^{T}$ | NZ_JRGK00000000 |
| *Aeromonas fluvialis* LMG 24681$^{T}$ | NZ_CDBO00000000 |
| *Aeromonas hydrophila* ATCC 7966$^{T}$ | NC_008570 |
| *Aeromonas jandaei* CECT 4228$^{T}$ | NZ_CDBV00000000 |
| *Aeromonas lacus* AE122$^{T}$ | NZ_JRGM00000000 |
| *Aeromonas media* CECT 4232$^{T}$ | NZ_CDBZ00000000.1 |
| *Aeromonas molluscorum* 848$^{T}$ | NZ_AQGQ00000000 |
| *Aeromonas piscicola* LMG 24783$^{T}$ | NZ_CDBL00000000 |
| *Aeromonas popoffii* CIP 105493$^{T}$ | NZ_CDBI00000000 |
| *Aeromonas rivipollensis* KN-Mc-11N1 | NZ_CP027856.1 |
| *Aeromonas rivuli* DSM 22539$^{T}$ | NZ_CDBJ00000000 |
| *Aeromonas salmonicida* subsp. *achromogenes* AS03 | NZ_AMQG00000000.2 |
| *Aeromonas salmonicida* subsp. *masoucida* NBRC 13784$^{T}$ | NZ_BAWQ00000000.1 |
| *Aeromonas salmonicida* subsp. *pectinolytica* 34mel$^{T}$ | ARYZ00000000.2 |
| *Aeromonas salmonicida* subsp. *salmonicida* ATCC 33658$^{T}$ | NZ_CDDW00000000.1 |
| *Aeromonas salmonicida* subsp. *smithia* JF4097 | NZ_JZTI00000000.1 |
| *Aeromonas sanarellii* LMG 24682$^{T}$ | NZ_CDBN00000000 |
| *Aeromonas schubertii* WL1483 | NZ_CP013067 |
| *Aeromonas simiae* CIP 107798$^{T}$ | NZ_CDBY00000000 |
| *Aeromonas sobria* CECT 4245$^{T}$ | CDBW00000000 |
| *Aeromonas taiwanensis* LMG 24683$^{T}$ | NZ_BAWK00000000 |
| *Aeromonas tecta* CECT 7082$^{T}$ | NZ_CDCA00000000 |
| *Aeromonas veronii* B565 | NC_015424 |
| *Escherichia coli* K-12 substr. BW25113 | NZ_CP009273.1 |
| *Pseudomonas aeruginosa* PAO1 | NC_002516.2 |
| *Pseudomonas extremaustralis* DSM 25547 | AHIP00000000.1 |
| *Pseudomonas protegens* Pf-5 | CP000076.1 |
| *Pseudomonas putida* KT2440 | NC_002947.4 |
| *Pseudomonas syringae* B728a | NC_007005.1 |
| *Shewanella oneidensis* MR-1$^{T}$ | NC_004347.2 |
